# HIF1α-dependent induction of the T-Type calcium channel CaV3.2 mediates hypoxia-induced neuronal hyperexcitability

**DOI:** 10.1101/2025.10.13.682038

**Authors:** Anna R. Tröscher, Despina Tsortouktzidis, Franziska Ammer-Pickhardt, Manuel Penzenleitner, Martin Aichholzer, Philip-Rudolf Rauch, Tobias Rossmann, Nico Stroh-Holly, Baran Alti, Andreas Gruber, Raimund Helbok, Jan Haubold, Christian Thome, Maren Engelhardt, Tim J. von Oertzen, Susanne Schoch, Albert J. Becker, Karen M.J. van Loo

## Abstract

Post-stroke epilepsy (PSE) is a leading cause of acquired epilepsy in adults, yet the molecular mechanisms linking post-ischemic hypoxia to neuronal hyperexcitability remain poorly understood. The transcription factor hypoxia-inducible factor 1α (HIF1α) is a central mediator of the cellular response to hypoxia and may contribute to epileptogenesis by regulating ion channel expression. Here, we identify the T-type calcium channel CaV3.2 (encoded by *Cacna1h*) as a direct transcriptional target of HIF1α and demonstrate its role in hypoxia-induced network hyperexcitability. Oxygen-glucose deprivation followed by reoxygenation (OGD/R) induced a persistent increase in neuronal firing rate. In murine and human organotypic brain slices, hypoxia led to a marked increase in *HIF1α* and *Cacna1h* expression at both transcript and protein levels. Using neuronal cell lines and primary cortical neurons we show that HIF1α activation through HIF1α overexpression, consistently increases *Cacna1h* expression. In NS20Y cells, overexpression of a normoxia-stable HIF1α variant increased *Cacna1h* promoter activity in both fluorescent and dual-luciferase reporter assays. The same effect was observed in primary cortical neurons, where HIF1α overexpression also elevated network activity measured by multielectrode array recordings, replicating the effect of the OGD/R model. Together, these results establish HIF1α as a key transcriptional regulator of CaV3.2 in neurons and reveal a conserved hypoxia-HIF1α-CaV3.2 pathway that enhances neuronal excitability. This mechanism may underlie hypoxia-induced network hyperactivity and contribute to the pathogenesis of post-ischemic epileptogenesis, offering a potential molecular target for intervention.

## Introduction

Post-stroke epilepsy (PSE) is a common complication among stroke survivors and accounts for a substantial proportion of acquired epilepsy cases in adults. In contrast to early seizures, which occur due to metabolic imbalances and blood-brain-barrier disruptions during the acute phase after stroke (Samake et al., 2023), PSE is characterized by the occurrence of unprovoked seizures more than seven days after a stroke, affecting approximately 20% of patients following severe ischemic stroke (Beghi et al., 2010; Galovic et al., 2018; Tanaka et al., 2024). Despite its clinical significance, the molecular mechanisms underlying hypoxia-induced epileptogenesis remain poorly understood.

Multiple processes contribute to PSE development, including permanent structural alterations, such as gliotic scarring, neurovascular dysfunction, maladaptive plasticity, and increased neuroinflammation (Galovic et al., 2018; Tröscher et al., 2021; Gruber et al., 2023). In addition, dynamic transcriptional alterations may drive the transition to an epileptogenic state. For example, the transcription factor early growth response protein 1 (Egr1) has been identified as a potential “master switch” of epileptogenesis, promoting neuronal hyperexcitability through the induction of channelopathies such as enhanced T-type calcium channel activity (Becker et al., 2008; van Loo et al., 2012, 2019).

The transcription factor hypoxia-inducible factor 1α (HIF1α) is a key mediator of the cellular response to hypoxia. Under normoxic conditions, HIF1α is continuously produced but rapidly degraded; during hypoxia, inhibition of hydroxylation stabilizes HIF1α, allowing it to dimerize with HIF1β and activate transcription of genes involved in cell death, energy metabolism, and angiogenesis (Sharp and Bernaudin, 2004). HIF1α may also contribute to post-ischemic epileptogenesis by directly regulating ion channel expression. In particular, it may bind to promoter regions of T-type calcium channel genes, initiating an epileptogenic cascade within neurons of the ischemic penumbra (Sellak et al., 2014). Among these channels, CaV3.2 has been repeatedly implicated in chronic epilepsy models (Becker et al., 2008; van Loo et al., 2012, 2019). It is upregulated up to three-fold during the early latent phase of epileptogenesis, leading to increased intrinsic bursting in hippocampal neurons (Sanabria et al., 2001; van Loo et al., 2012).

Outside the brain, hypoxia-induced upregulation of CaV3.2 has been demonstrated in glomus cells, PC12 cells, and cardiomyocytes (Del Toro et al., 2003; Makarenko et al., 2016; Morishima et al., 2021), as well as in pulmonary smooth muscle cells where HIF1α-mediated CaV3.2 expression enhances T-type channel currents (Sellak et al., 2014).

To date, the role of HIF1α in regulating CaV3.2 expression in the brain, and its potential contribution to epileptogenesis, remain unknown. Given these observations, we hypothesized that hypoxia-induced HIF1α activation increases CaV3.2 expression, thereby enhancing neuronal excitability. To address this, we investigated the relationship between HIF1α activation and CaV3.2 expression in NS20Y cells, primary neuronal cultures, and organotypic brain slice cultures from both human and mouse tissue. We demonstrate that CaV3.2 expression is increased following hypoxia as well as after HIF1α overexpression alone. Moreover, multiple electrode array recordings revealed a corresponding increase in network excitability, reflected by a higher weighted mean firing rate upon HIF1α activation.

## Materials and Methods

### Neuronal Cultures

All animals were handled according to government regulations as approved by local authorities (LANUV Recklinghausen) and all procedures were performed according to the guidelines of the University Hospital Bonn Animal-Care-Committee as well as the European Directive (2010/63&EU).

Pregnant C57BL/6 mice were deeply anesthetized at embryonic day 16 and sacrificed by decapitation. Abdominal skin and uterus were opened, embryos extracted and kept in PBS (Gibco #14190094) on ice. After decapitation of embryos, brains were removed and divided in sagittal fashion, meninges and cerebellum were removed, and cortex was dissected and placed in cold HBSS (Gibco #14170138). Tissue was digested with Trypsin (Gibco #15090046, 1:10 in HBSS) for 20 min. Next, cells were dissociated with DNase I (Roche #112484932001) and BME (Gibco #41010109), supplemented with FBS. Cell culture plates were coated for 1h at room temperature with Poly-D-lysine (Sigma-Aldrich #11284932001). 70.000 Cells were seeded per well in a 24-well plate. Cells were kept in BME medium (Gibco #21010046), supplemented with 1 % FBS, 0.5 mM L-Glutamine, 0.5 % glucose (Gibco #15384895) and incubated at 37 °C, 5% CO_2_. For multielectrode array (MEA) experiments, 40.000 neurons were seeded per well in a 24-well Cytoview-MEA plate (Axion #M384-tMEA-24W).

### Oxygen-glucose deprivation/re-perfusion model (OGD/R)

Primary neuronal cultures were grown under standard conditions until days in vitro (DIV) 11/12 to enable neuronal maturation and network formation. For OGD/R, HBSS was bubbled with a 95 % N_2_/5 % CO_2_ gas for 5 min with 8-10L/min to deoxygenate the incubation medium. Standard neuronal culturing medium was removed and collected. Normoxic or deoxygenated HBSS were added to control and OGD/R cultures, respectively. OGD/R cultures were placed in the hypoxia incubation chamber (Stemmcell technologies, #27310) and the chamber was filled with a 95 % N_2_/5 % CO_2_ gas for 4 min with a flowrate of 20L/min. All cells were incubated for 15 min in the incubator, then OGD/R cultures were removed from the chamber and HBSS was removed from all cultures and replaced with the original culturing medium.

### MEA Recordings

Between DIV13/14 to DIV 22/24 of primary neuronal cultures, electrical activity of the cultures was measured using the Maestro Edge MEA system of Axion Biosystems every two days in 24-well plates including 16 electrodes per well (4×4 grid, recording area: 1.1mmx1.1mm, electrode spacing: 350 µm). Plates were kept at 37 °C and 5 % CO_2_ during the whole time of the recording. Plates were put in the apparatus, left to equilibrate for 10 min, followed by 10 min recording of spontaneous activity with the Axis Navigator software (version 2.0.4.21). Raw recording data were analyzed with the same software with the following settings: Spike detection: 6 x SD from voltage baseline, minimal burst criterion: minimum of 5 spikes, maximum inter-spike interval: 100 ms. Spikes traces were generated by the Axion Neural Metric Tool (version 3.2.25) and voltage traces using the Axion Data Export Tool (version 3.4.5.). For analyses, only wells with at least 15 active electrodes at the first measurement were used. The weighted mean firing rate was calculated (mean firing rate per well, corrected for the number of active electrodes) and normalized for each well separately to the activity at the first measurement. For analysis, measurements from DIV13-14 (Bin 1), DIV15-17 (Bin 2), DIV 18/19 (Bin 3) and DIV22-24 (Bin 4) were pooled.

### Mouse organotypic brain slice cultures

Nine week old C57BL/J6 mice (males/females) were euthanized by cervical dislocation and brains were quickly extracted. The whole brain was cut coronally in 300 µm sections on a vibratome in oxygenated aCSF (124mM NaCl, 26 mM NaHCO_3_, 10mM D-Glucose, 3 mM KCl, 1.25 mM NaH_2_PO_4_, 2 mM CaCl_2_, 1 mM MgSO_4_, 10 mM HEPES). Brain sections were split into single hemispheres and put on filter membranes (BRAND®, #782720) in 12-well plates and incubated with organotypic slice medium (66 % Neurobasal-A (Gibco #10888-022), 33 % HBSS (Gibco, #14025-092), 0.625 % D-Glucose (Gibco, #A2494001), 0.01 M HEPES (Gibco, #15630049), 2 mM L-Glutamine (Gibco, #25030-081), 0.5 X B27 (Gibco, #A35828-01), 2 % Anti-Anti (Merck, #A5955) at 35 °C and 5 % CO_2_. One hemisphere was used for normoxic control and the other hemisphere for hypoxia. On DIV3, half the slices were put in a hypoxia cell culture system (BioSpherix X3 Xvivo System) with 3 % O_2_. After four days in hypoxic conditions, hypoxic and normoxic samples were used for RNA isolation.

### RNA isolation, cDNA synthesis and qPCR analysis

RNA from mouse OTCs was isolated with the RNeasy mini kit (Qiagen, #74104) according to the manufacturer’s instructions. Subsequently, 40 ng of RNA was reverse transcribed using the Luna RT Master mix (New Englad Biolab, #E3010L). qPCR was performed with the Luna Universal qPCR master mix (New England Biolabs, #M300S) according to the manufacturer’s instructions. In short, a 20 µl PCR reaction was used for each sample, including 0.25 µM forward and reverse primers and 1 µl cDNA. Cycling was performed with a 60 sec initial denaturation, followed by 40 rounds of 15 seconds at 95 °C and 45 seconds at 60 °C. Standard melting curve was performed after qPCR amplification. Synaptophysin was used as housekeeping gene for normalization. Primers used were the following: Synaptophysin: 5’-TATCAACCCGATTACGGGCA-3’ and 5’-TGGGCTTCACTGACCAGATT-3’; HIF1-alpha: 5’-CCATTTTCAACTCAGGACACTG-3’ and 5’-GTGCTCATACTTGGAGGGCT-3’; CaV3.2: 5’-ATGTCATCACCATGTCCATGGA-3’ and 5’-ACGTAGTTGCAGTACTTAAGGGCC-3’.

### Human organotypic brain slice cultures

All patients gave informed written consent and the study was approved by the Ethics commission of the Johannes Kepler University, under the number EK1250/2023. Patient details can be found in Supplementary Table 1. After surgical resection, human tissue from the access path of tumor resections or epilepsy surgery was quickly transported to the lab in aCSF, where blood vessels and meninges were removed, before it was cut into 300 µm sections in oxygenated aCSF at the vibratome. Sections were transferred to filter membranes in 6-well plates and incubated in organotypic slice medium at 35 °C and 5 % CO_2_. At DIV3, half of the human tissue was transferred to the oxygen-controlled incubator and incubated at 3 % O_2_ for 4 days (until DIV7). From each patient, one section from each condition was directly fixed for 90 min in 2 % PFA and another section was used for western blot analysis.

### Western Blot analysis

Tissue was lysed in RIPA buffer substituted with Halt Protease and Phosphatase Inhibitor Cocktail (100X, Thermo Scientific #78440). For each 1 mg tissue 10 µl buffer was used and tissue was subjected to 3 cycles of freezing and thawing to break up cell membranes. The amount of protein isolated was quantified with the Pierce BCA Protein assay (Thermo Scientific #A65453) and 20 ug were loaded on a 10 % SDS-PAGE gel. Proteins were transferred to a nitrocellulose membrane (Bio-Rad #1620115) and Ponceau S (ThermoFisher #A4000279) staining was performed to visualize total protein content. After imaging of the Ponceau S-stained membranes and destaining, unspecific antibody binding was blocked by 2% fish block (2% fish skin gelatin (Sigma #G7041-100G) in PBS-T). Primary antibodies against HIF1α (Proteintech #20960-1-AP) were used in a concentration of 1:1000 in fish block over night at 4 °C. Biotinylated secondary antibodies (Goat anti-Rabbit IgG (h+L) Cross-Absorbed Secondary Antibody, HRP, #G-21234, 1:5000) were used for 45 min at room temperature. The membrane was then incubated for 5 min in the Immobilon Western HRP Substrate (Millipore #WBKLS0500) and imaged at the Bio-Rad ChemiDoc MP Imaging System. Quantification of bands was performed with the Image Lab software (Image Lab 6.1) from Biorad and whole protein normalization was used.

### Immunofluorescence Staining and Imaging

Human OTCs, which showed an upregulation of HIF1α in the western blot analysis, were used for free floating immunofluorescence stainings. To this end, fixed sections were incubated in fish block (1 % BSA. 0.2 % fish skin gelatin (Sigma #G7041-100G), 0.01 % triton X-100(Merck #102419574) for 1 h at room temperature, followed by overnight incubation at 4 °C with primary antibodies against NeuN (1:1000, Merck #4182374) and CaV3.2 (1:1000, Invitrogen #8M1106). Secondary antibody incubation was performed with an anti-chicken 647 (1:5000, Invitrogen #2335730) and an anti-rabbit 488 (1:5000, Invitrogen #A-11001) antibody. Sections were mounted on glass slides, covered in ROTI Mount FluorCare (Carl Roth # HP19.1). Sections were imaged at a confocal microscope (Leica Stellaris 5) with the same intensity settings for each paired sample (normoxic and hypoxic). For the quantification of CaV3.2 content, the percentage of the image covered by the positive CaV3.2 signal, using the same threshold settings for each matched sample, was calculated using Fiji software.

### Constructs

The pAAV-syn-mHif1a-T2A-EGFP plasmid was generated by a two fragment in-fusion cloning (FC; Takara Bio Europe/Clontech). pcDNA3-mHIF-1α-MYC (P402A/P577A/N813A) (Addgene #44028, (Hu et al., 2007)), a plasmid with a normoxic stable variant of HIF1 α, was used as a PCR template for the mHIF1α fragment (Forward primer: 5’-cgcatcgattgaattatggagggcgccgg-3’, reverse primer: 5’-tcgctagccagatcctcttctgagatgagtttttgtt-3’), and the pLenti-U6-sgRNALacZ-Syn-dCas9-KRAB-T2A-eGFP plasmid (Tsortouktzidis et al., 2022) was used as a PCR template for the T2A-EGFP fragment (forward primer: 5’-ggatctggctagcgagggcagagg-3’, reverse primer: 5’-ccgctcggtccgcacgcggggaggcggc-3’). Both fragments were subsequently inserted in the *EcoR*I/*Pml*I sites of the pAAV-hSyn-MCS plasmid (Tsortouktzidis et al., 2022).

### Cell Culture, Transfection and Luciferase Assay

NS20Y cells, a murine cholinergic neuroblastoma cell line (Sigma, #08062517), were maintained at 37 °C, 5 % CO_2_ in DMEM (Sigma, D6546) supplemented with 10 % heat inactivated FBS (Gibco #10270106), 2 mM L-Glutamine (Gibco #11539876) and 100 units/mL penicillin/streptomycin (Gibco #15140122). Cells were transfected in 24-well plates using Lipofectamine (Invitrogen, #11668030) with the following DNA concentrations per well: pAAV-hSyn-HIF1 300 ng or CAG-GFP 300 ng in controls, Cacna1h-Luciferase 100 ng and Renilla 25 ng (luciferase experiments), or Cacna1h-mRuby 200 ng and CAG-GFP 75 ng (imaging experiments). 48 h after transfection, cells were imaged or collected for luciferase assays. Dual luciferase reporter assays (Promega, #E1910) were performed according to the manufacturers’ instructions and firefly/Renilla luciferase ratios were determined using the Glomax Luminometer (Promega).

### Cell Imaging

Images were obtained using an inverted phase contrast fluorescent microscope (Zeiss Axio Observer A1 with objectives 20X, LD A-Plan) and processed using Fiji. The cell surface area, set by GFP fluorescent signal, was determined using the Weka trainable segmentation plugin. Integrated density, measured by mRuby fluorescent signal, was normalized to the cell surface area to obtain reported values of integrated density/cell surface.

### Viral Production and Neuronal Transduction

AAV1/2 viruses were produced in HEK293-AAV cells (Agilent, #240073) by triple CaPO_4_ transfection as described previously (van Loo et al., 2012). Transduction of primary neurons was performed at days DIV4 in 24 well plates. For luciferase analyses cells were collected at DIV12-13.

### Artificial Neural Network Analysis of MEA Data

We developed an artificial neural network to compare neuronal network dynamics across experimental conditions in an unsupervised manner. The classifier was trained on multielectrode array (MEA) recordings from primary cortical neurons. The goal of this analysis was to determine how closely the activity patterns of HIF1α-overexpressing (HIF^+^) and control (GFP) cultures under normoxia resembled those of wild-type cultures maintained under hypoxic or normoxic conditions. Raw extracellular voltage traces from each MEA well (16 electrodes per well) were first processed to remove noise and then partitioned by culture. Each culture (one MEA well, 16 electrodes) was treated as an independent sample to prevent temporal or spatial data leakage between datasets. Wells from wild-type hypoxic and wild-type normoxic conditions were randomly divided into training (≈33%), validation (≈33%), and test (≈33%) subsets. Only wild-type conditions were used for model optimization and performance evaluation. The entirely separate set of HIF^+^ and GFP cultures, recorded under normoxia, served exclusively as an external test cohort for biological interpretation of network behavior. Each MEA recording was segmented into 100 ms time windows, which were independently classified by the network. To account for developmental changes in culture maturation, separate CNN– LSTM models were trained for each recording time bin (DIV13–14, DIV15–17, DIV18–19, DIV22–24). The CNN-LSTM architecture combined a convolutional neural network (CNN) to extract relevant features and a long short-term memory (LSTM) network to capture temporal dependencies. Model outputs were continuous probabilities (0–1) corresponding to “hypoxia-like” or “normoxia-like” network activity. Generalization was improved by blass reweighting, weight decay, and learning rate scheduling. Only data from independent electrodes (well excluded from training) were used for plotting and downstream interpretation. Further details on model architecture are provided in the Supplementary Materials.

### Statistics

Two-way repeated measures ANOVA with Sidaks multiple comparison correction was performed for analysis of MEA recordings (weighted mean firing rate: WMFR) over time. Results from qPCR, western blot and immunofluorescent stainings were analyzed with paired T tests. Analysis of fluorescent reporter activity in NS20Y cells was performed by unpaired T tests. Luciferase dual reporter assay analyses were analyzed using one-sample T Tests. An alpha value of below 0.05 was considered to be statistically significant.

## Results

### Hypoxia increases neuronal firing rate

We first investigated the functional consequences of hypoxia on neuronal network activity. MEA recordings were performed on primary cortical neurons maintained under normoxia or exposed to 15 min of hypoxia followed by reoxygenation (OGD/R). Cultures subjected to OGD/R exhibited increased neuronal firing compared with normoxic controls in representative recordings at DIV18 (Fig 1A, B). Over the subsequent 10 days, normoxic control cultures showed a mild, maturation-related increase in WMFR, whereas OGD/R-treated cultures exhibited a significantly greater and sustained increase (two-way ANOVA hypoxia × normoxia: F = 7.66, p = 0.009; Fig. 1C).

**Fig 1.**
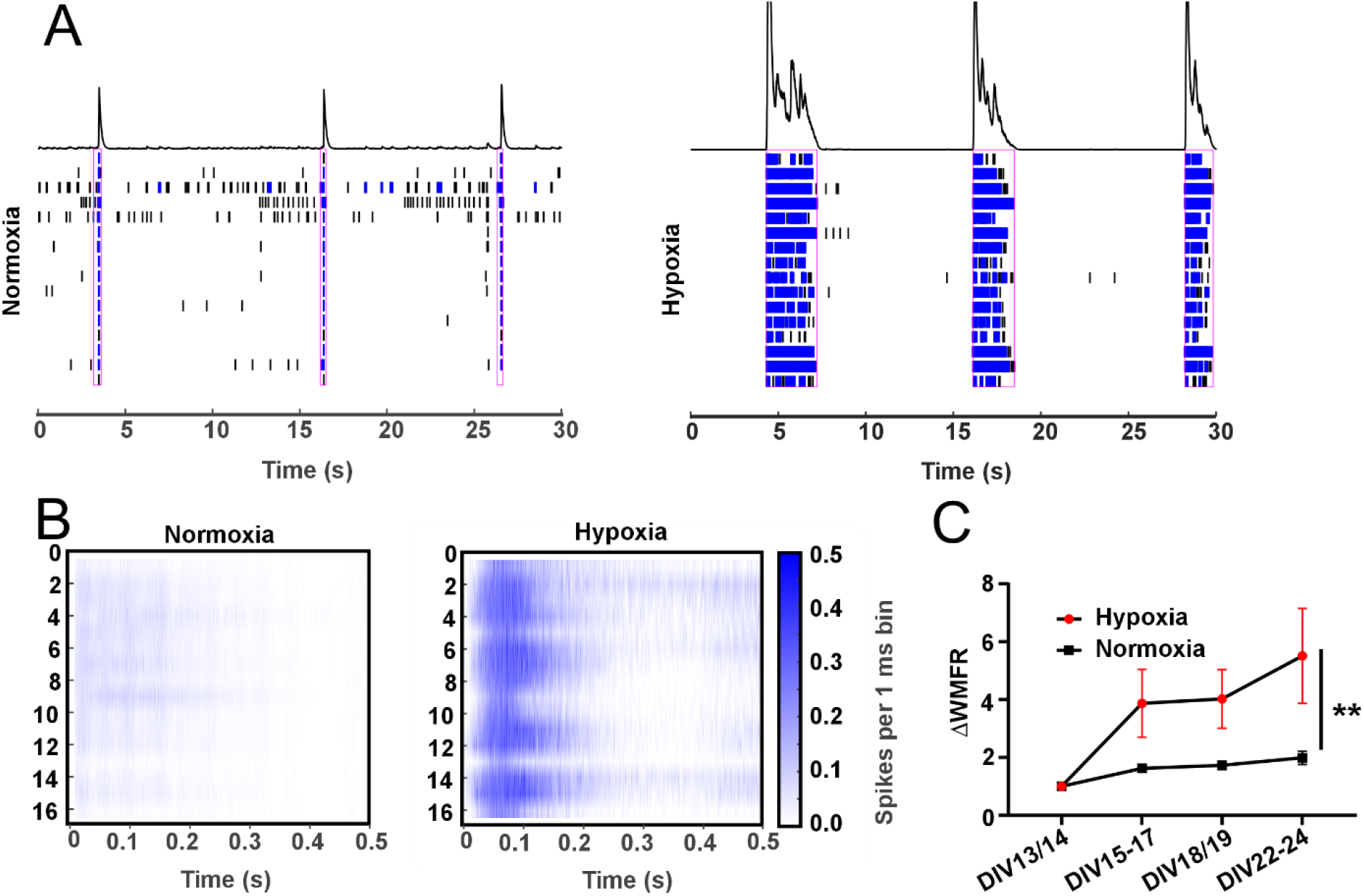
OGD/R leads to increased neural firing rate. Representative images of (A) network bursts and (B) activity recorded from every electrode in normoxic controls and OGD/R-treated neurons from primary cortical cultures at DIV18. (C) Primary neurons exposed to OGD/R showed a significantly increased weighted mean firing rate (WMFR) compared to neurons maintained under normoxic conditions throughout the experiment (n = 15 and 25 wells, respectively; two-way ANOVA hypoxia × normoxia: F = 7.66, p = 0.009).

### Hypoxia enhances HIF1α and CaV3.2 expression in organotypic brain slice cultures

To extend these findings to a more physiologically relevant system, we analyzed murine OTCs exposed to hypoxia or maintained under normoxic conditions. Hypoxia led to a fourfold increase in *Hif1a* expression, confirming successful induction of a hypoxic cellular response (p = 0.05; Fig. 2A). In parallel, CaV3.2 expression increased 3.5-fold compared with normoxic controls (p = 0.009; Fig. 2B), indicating that hypoxia enhances both Hif1α and CaV3.2 expression in brain tissue.

**Fig 2.**
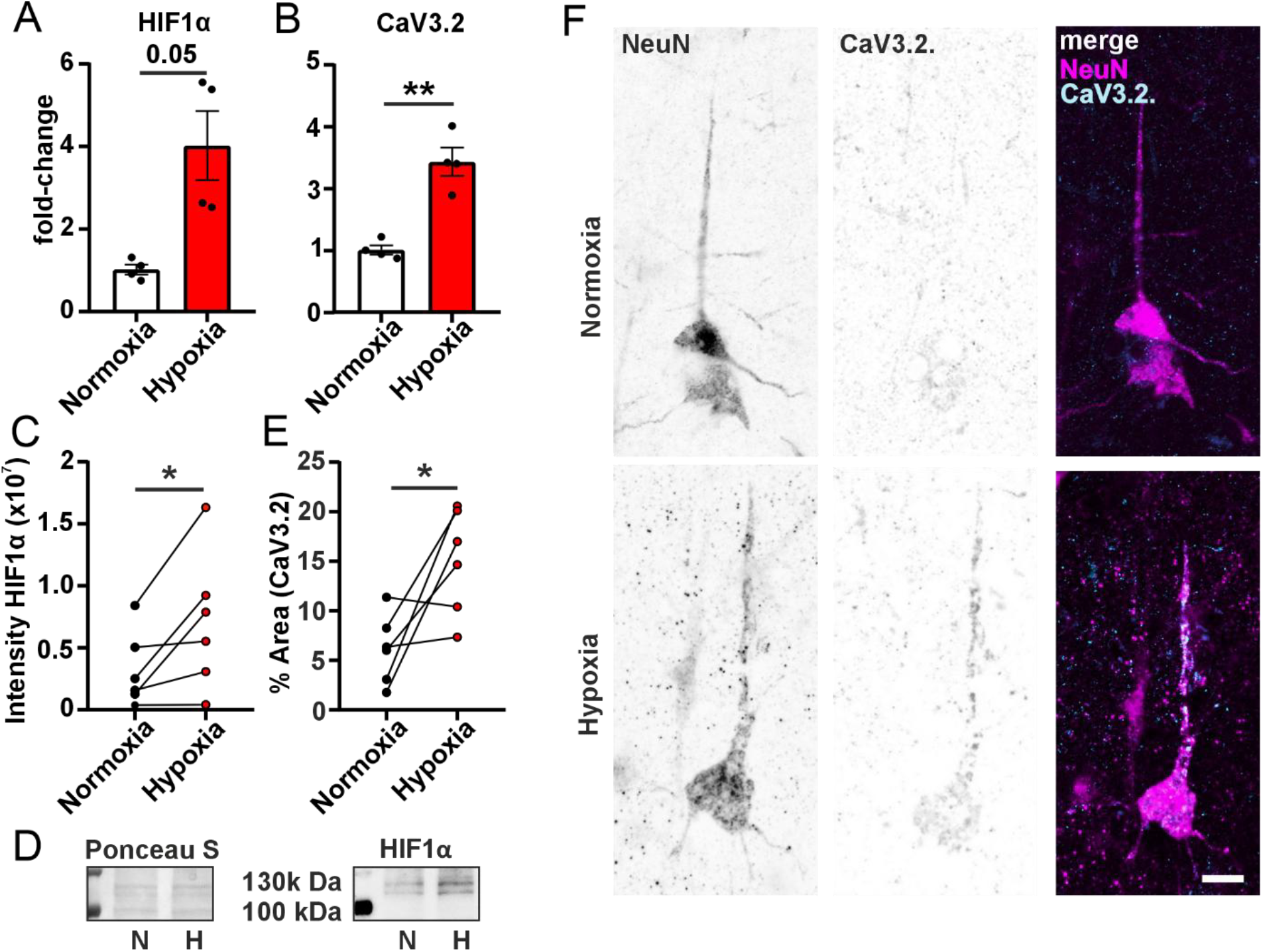
Upregulation of CaV3.2 after hypoxia in organotypic brain slice cultures (OTCs). (A) *Hif1α* mRNA was significantly increased in murine OTCs following hypoxia compared with normoxic controls (two-sided paired T-Test, p = 0.05). (B) *Cacna1h* expression showed a strong increase upon hypoxia (two-sided paired T-Test, **p = 0.009). (C) Human OTCs exposed to hypoxia exhibited significantly increased HIF1α protein levels by western blot analysis (two-sided paired T-test, *p = 0.04). (D) Representative HIF1α bands (at 120 kDa) and Ponceau S staining of normoxia and hypoxia OTCs. (E) Quantification of immunofluorescence revealed a significantly larger CaV3.2-positive area under hypoxia compared to normoxia (two-sided paired T-Test, *p = 0.03). (F) Representative images of NeuN-positive cells (neurons) and CaV3.2 staining illustrate low CaV3.2 baseline expression under normoxia and pronounced upregulation along apical dendrites after hypoxia. Scale bar: 10 µm.

We next assessed whether this response is conserved in human OTCs. Western blot analysis confirmed a significant increase in HIF1α protein levels under hypoxic conditions (p = 0.04; Fig. 2C, D). Immunofluorescence staining further revealed a significant expansion of the CaV3.2-positive area following hypoxia (p = 0.03; Fig. 2E). Representative images show enhanced CaV3.2 immunoreactivity in dendrites of NeuN-positive neurons in hypoxia-treated slices compared with normoxic controls (Fig. 2F).

Together, these results demonstrate that hypoxia induces HIF1α stabilization and CaV3.2 upregulation in both murine and human OTC, indicating a conserved mechanism by which hypoxic signaling enhances T-type calcium channel expression in neurons.

### HIF1α regulates CaV3.2 expression in neuronal cell lines

To determine whether HIF1α directly regulates expression of the T-type calcium channel CaV3.2 and contributes to hypoxia-induced neuronal hyperexcitability, we first performed an *in silico* analysis of the human *Cacna1h* promoter region. Using the JASPAR database with a relative profile score threshold of 80%, we screened the 2,135 nucleotides upstream of the transcriptional start site (ATG) for potential HIF1α binding motifs. This analysis revealed 44 putative HIF1α binding sites within the *Cacna1h* promoter region (Supplementary Table 2), suggesting that *Cacna1h* may be a direct transcriptional target of HIF1α.

To test this experimentally, we transfected the neuronal cell line NS20Y with either a control plasmid (CAG–GFP) or a plasmid encoding a normoxia-stable HIF1α variant (hSyn–HIF1α–GFP), together with a fluorescent reporter driven by the *Cacna1h* promoter (Tsortouktzidis et al., 2022). Cells expressing HIF1α displayed a significant increase in mRuby reporter fluorescence, indicating enhanced *Cacna1h* promoter activity (p = 0.013; Fig. 3A).

**Fig 3.**
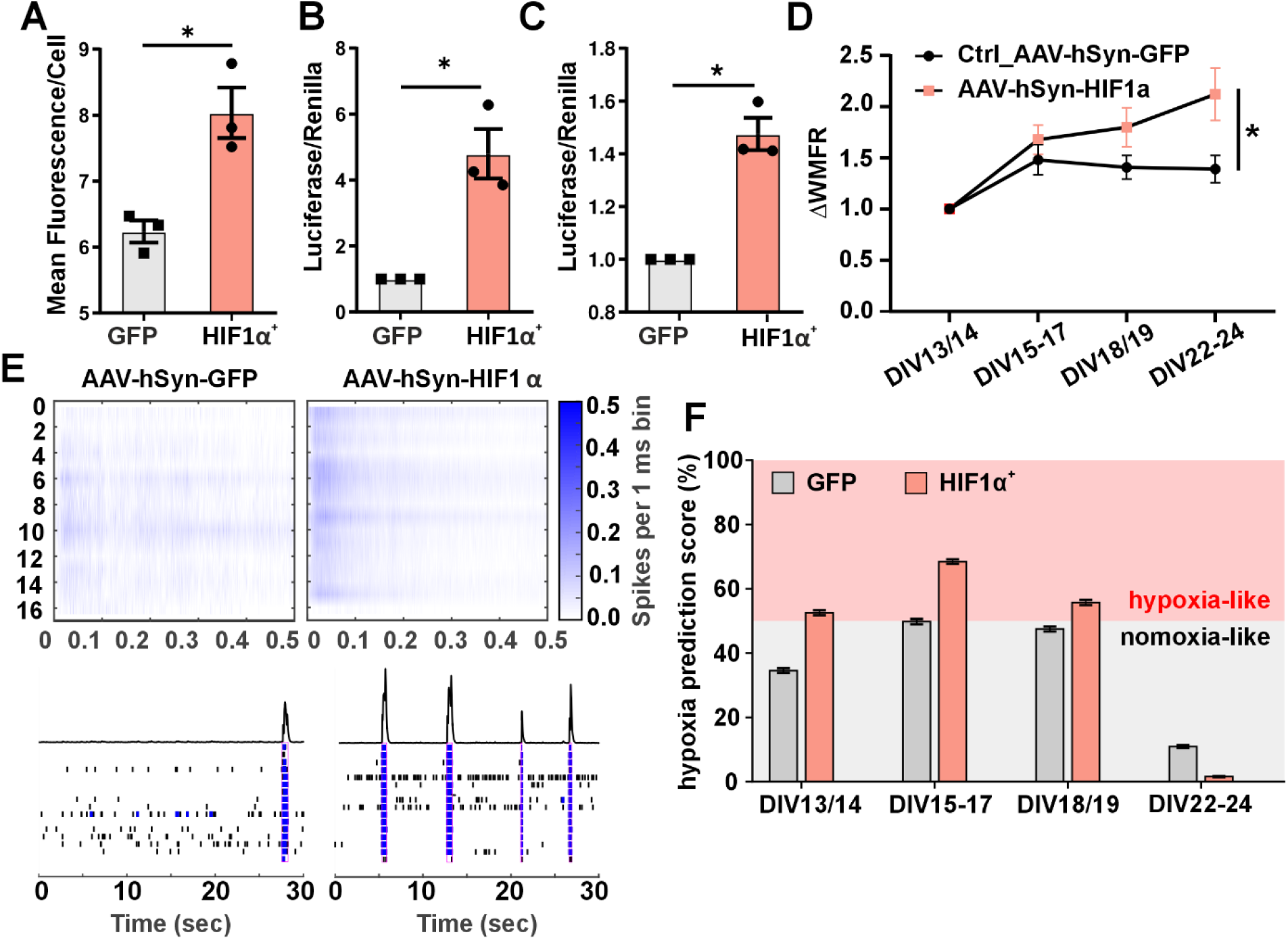
Upregulation of CaV3.2 upon HIF1α transduction induces increased weighted mean firing rate (WMFR). NS20Y cells transfected with HIF1α show a significant upregulation of CaV3.2 in (A) a fluorescent reporter assay (Two-sided T Test, *p = 0.013), (B) as well as in a dual-luciferase reporter assay (3 biological replicates, one-sample T Test, *p = 0.037), and (C) primary cortical neurons (3 biological replicates, one-sample T-Test, *p = 0.016). (D) Primary neurons transduced with AAV-hSyn-HIF1α (n = 18 wells) show significantly increased WMFR compared to primary neurons transduced with a control virus (n = 18 wells; AAV-hSyn-GFP, two-way ANOVA HIF1α^+^/GFP: F = 4.17, *p = 0.049). (E) Representative recordings of AAV-hSyn-GFP and AAV-hSyn-HIF1α cultures at DIV18 show electrode activity (upper panels) and bursting behavior (lower panels). (F) Convolutional neural network–long short-term memory (CNN–LSTM) classifier analysis confirmed more hypoxia-like network behavior in HIF1α-transduced neurons compared to controls.

To validate these findings, we repeated the experiment using a dual-luciferase reporter assay (Tsortouktzidis et al., 2022). HIF1α overexpression again significantly increased *Cacna1h* promoter-driven luciferase activity compared with controls (p = 0.037; Fig. 3B). These results demonstrate that HIF1α is sufficient to activate *Cacna1h* promoter activation in neuronal cell lines.

### HIF1α induces CaV3.2 promoter activity and increases excitability in primary neurons

We next examined whether this regulatory relationship is maintained in primary neurons. Murine primary cortical cultures were co-transfected with the dual-luciferase reporter system and either a control plasmid or the HIF1α construct. Consistent with the results in NS20Y cells, HIF1α expression significantly increased *Cacna1h* promoter activity, serving as a surrogate for CaV3.2 mRNA expression in primary neurons (p = 0.016; Fig. 3C).

To determine whether this transcriptional activation translates into altered neuronal activity, we assessed network dynamics using MEA recordings following adeno-associated viral (AAV) transduction with either control (rAAV-hSyn–GFP) or HIF1α (rAAV-hSyn–HIF1α–GFP) vectors. HIF1α overexpression led to a significant increase in the WMFR compared with controls (two-way ANOVA HIF1α^+^/GFP: F = 4.17, p = 0.049; Fig. 3D, E).

We then applied a previously trained convolutional neural network–long short-term memory (CNN– LSTM) classifier, developed to distinguish hypoxic from normoxic network activity, to this dataset. Without retraining or parameter adjustment, the network categorized most HIF1α-overexpressing samples as displaying “hypoxia-like” dynamics, while controls were classified as “normoxia-like,” except for bin 4 (Fig. 3F). This independent analysis confirmed that HIF1α overexpression alone reproduces activity patterns characteristic of hypoxia-induced hyperactivity.

Across complementary experimental systems, from neuronal cell lines to primary neurons and organotypic brain slice cultures, HIF1α activation consistently upregulated CaV3.2 expression and enhanced neuronal firing. These data identify HIF1α as a key transcriptional regulator of *Cacna1h* and suggest that HIF1α-driven CaV3.2 induction contributes to hypoxia-induced network hyperexcitability relevant to post-ischemic epileptogenesis.

## Discussion

PSE is a severe and frequent complication of ischemic stroke, affecting up to 20% of patients and currently lacking preventive treatment options (Beghi et al., 2010; Galovic et al., 2018; Tanaka et al., 2024). At present, patients are treated only after the first seizure, emphasizing the urgent need to identify molecular mechanisms that could be targeted to prevent epileptogenesis in high-risk individuals. PSE is a multifactorial disorder involving gliotic scar formation, chronic neuroinflammation,and neurovascular dysfunction (Galovic et al., 2018; Tröscher et al., 2021; Gruber et al., 2023). However, the cellular and molecular changes that precede network hyperexcitability remain poorly defined.

Here, we identify HIF1α as a transcriptional regulator of the T-type calcium channel CaV3.2 and demonstrate that this pathway contributes to hypoxia-induced neuronal hyperexcitability. Using complementary models, including NS20Y cells, primary neuronal cultures, and OTCs from mouse and human, we show that activation of HIF1α, either by transient hypoxia or direct overexpression, leads to *Cacna1h* upregulation and increased neuronal firing. These findings establish a direct molecular link between hypoxic signaling and the development of hyperexcitable neuronal networks, providing a potential mechanistic basis for post-ischemic epileptogenesis.

HIF1α is a central transcription factor orchestrating cellular adaptation to low oxygen by regulating genes involved in metabolism, angiogenesis, and survival (Sharp and Bernaudin, 2004). In neurons, accumulating evidence suggests additional roles beyond metabolic adaptation, including modulation of synaptic activity and ion channel expression (Wang et al., 2015; Arias-Cavieres et al., 2020; Zhang et al., 2023; Sobrido-Cameán et al., 2025). We show that *Cacna1h*, which encodes the T-type calcium channel CaV3.2, is transcriptionally activated by HIF1α. Using reporter assays, we show that HIF1α alone, under normoxic conditions, induces CaV3.2 expression in NS20Y cells and primary neurons. This finding aligns with bioinformatic predictions of HIF1α binding motifs within the *Cacna1h* promoter and reveals a previously unknown regulatory mechanism in the central nervous system.

Previous studies have shown that *CaV3*.*2* is transiently upregulated following pilocarpine-induced status epilepticus, where it contributes to epileptogenesis by promoting intrinsic bursting and network synchronization (Becker et al., 2008; van Loo et al., 2012, 2019). Similarly, increased *Cacna1h* expression has been reported in peripheral cell types exposed to hypoxia, including glomus cells and cardiomyocytes (Del Toro et al., 2003; Makarenko et al., 2016; Morishima et al., 2021). Our data demonstrate that this mechanism operates in neurons as well as in murine and human tissue, establishing a conserved hypoxia-responsive pathway.

Functionally, both hypoxia and HIF1α overexpression increased network activity over time. MEA recordings revealed that neurons exposed to OGD/R develop a persistent increased in WMFR lasting more than 10 days, consistent with long-term changes in excitability rather than transient metabolic effects. Therefore, this mechanism is less likely involved in acute symptomatic seizures, occurring within seven days after stroke, but rather parallels the time frame of the latent period before clinical seizure onset in PSE, defined as seizures occurring more than one week after stroke (Samake et al., 2023). Notably, HIF1α overexpression alone recapitulated the same sustained increase in WMFR, confirming that HIF1α activation is sufficient to induce electrophysiological changes characteristic of hypoxia-treated neurons. Artificial neural network analyses independently classified both HIF1α-overexpressing or hypoxia-treated cultures as “hyperexcitable” or “hypoxia-like.” Increased CaV3.2 expression likely contributes to these electrophysiological changes, as CaV3.2 channels mediate low-threshold calcium spikes that promote burst firing and oscillatory activity in cortical and thalamic circuits (Tringham et al., 2012; Cain et al., 2018; Dumenieu et al., 2018; Harding and Zamponi, 2022). Thus, HIF1α-driven upregulation of *Cacna1h* may facilitate calcium influx, promote activity-dependent transcription, and initiate long-term plasticity programs associated with epileptogenesis.

Our findings suggest that HIF1α–CaV3.2 signaling represents an early molecular event linking hypoxia to network hyperexcitability. In the ischemic penumbra, neurons experience reduced oxygen and glucose but remain viable, undergoing transcriptional remodeling that may predispose them to aberrant excitability (Stroemer et al., 1995; Ginsberg, 2003). Persistent HIF1α activation in these neurons could drive CaV3.2-dependent calcium entry and secondary signaling cascades, contributing to maladaptive plasticity and, ultimately, seizure generation.

From a clinical perspective, approximately 20 % of patients with severe stroke develop PSE (Galovic et al., 2018). This may indicate, that activation of the HIF1α–CaV3.2 axis occurs in vulnerable neuronal populations in certain brain regions or network nodes, which were shown to have a higher probability of being involved in PSE (Gruber et al., 2025). The risk of PSE correlates with the infarct size (Galovic et al., 2018), suggesting that widespread hypoxia-driven signaling could engage multiple connected regions. Notably, limiting hypoxia via thrombectomy in large-vessel occlusion does not reduce PSE risk (Gruber et al., 2023), consistent with our findings that even brief hypoxic episodes (15 minutes) activate the HIF1α-CaV3.2 pathway.

In summary, we identify a novel hypoxia–HIF1α–CaV3.2 signaling axis that drives neuronal hyperexcitability and may contribute to PSE. By demonstrating that HIF1α alone is sufficient to induce *Cacna1h* expression and increase neuronal firing, this study expands the known repertoire of HIF1α target genes and establishes CaV3.2 as a mechanistic link between hypoxia and long-term excitability changes. Because both HIF1α and CaV3.2 are pharmacologically targetable, this pathway may represent a promising therapeutic target to prevent or mitigate epileptogenesis following ischemic injury.

## Supporting information

Model Description

## Acknowledgements

We thank Pia Trebing, Sabine Opitz and Tobias Herbinger for technical assistance and Sabina Köfler for providing access to a high-performance computing server used for model training.

