## Supplementary figures and images for "HIF1α-dependent induction of the T-Type calcium channel CaV3.2 mediates hypoxia-induced neuronal hyperexcitability"

### Supplementary Figure 1

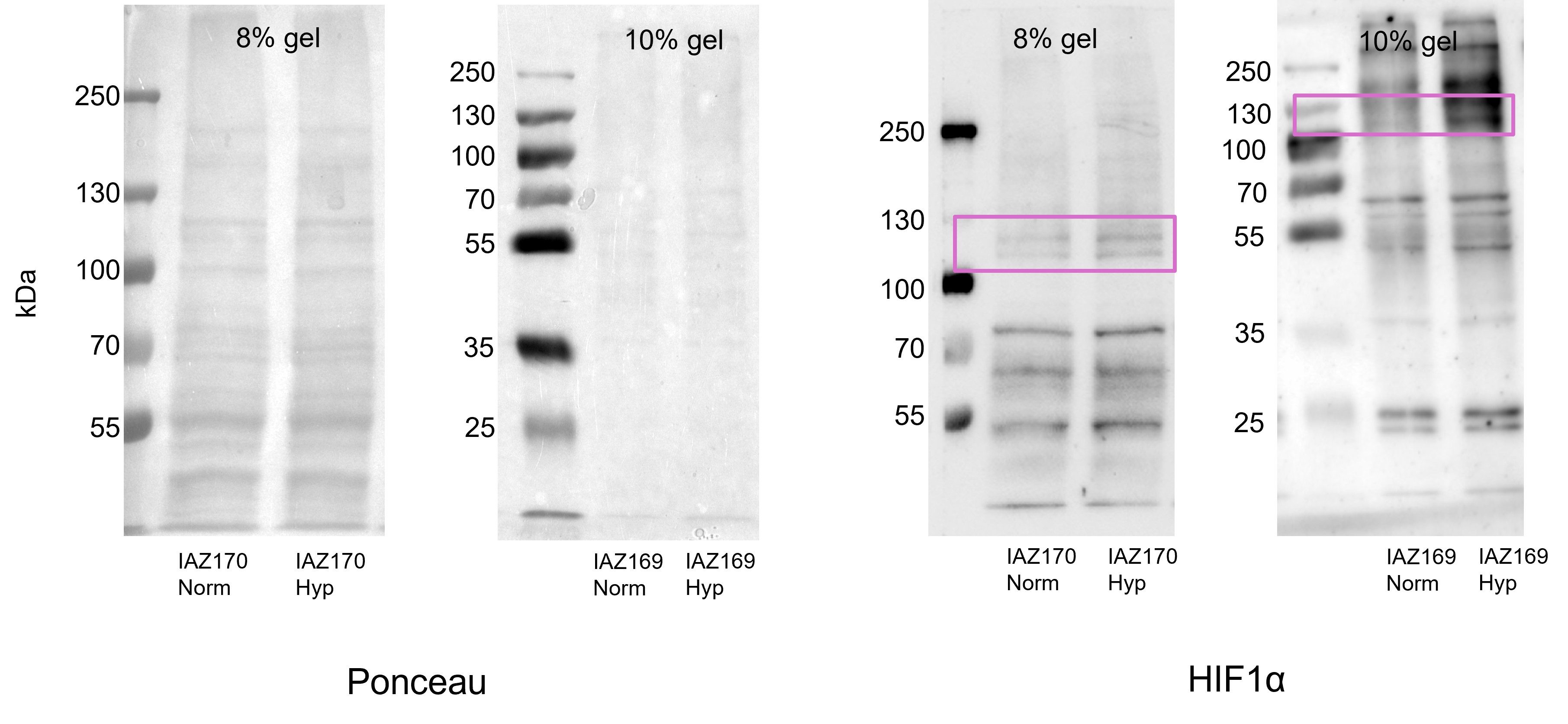
